# DECOUPLING OF EMIGRATION TIMING AND SKILL ACQUISITION IN A SATURATED POPULATION OF GOLDEN EAGLES

**DOI:** 10.1101/2025.04.21.649390

**Authors:** Hester Brønnvik, Martina Scacco, Michael Chimento, Enrico Bassi, Wolfgang Fiedler, Julia S. Hatzl, Petra Sumasgutner, Matthias Tschumi, Martin Wikelski, Svea Zimmermann, Kamran Safi, Elham Nourani

## Abstract

Many young animals must gain independence from parental care by acquiring the necessary skills for survival. Thus, the rate of skill acquisition can predict how long juveniles remain with their parents before emigrating from the natal territory. Yet in some systems, young animals acquire adult-like skills and the capacity to be independent long before they emigrate. This raises the question of why the timing of emigration decouples from the rate of skill acquisition, resulting in extended parental care. We addressed this by leveraging GPS tracking data from juvenile golden eagles (*Aquila chrysaetos*), which learn to fly and to hunt while remaining with their parents for 61-283 days after fledging. Young golden eagles often go on excursions without their parents outside of their natal territories before emigration. We expected flight performance to predict the timing of excursions better than the timing of emigration, which should be more influenced by external factors. In a set of time-to-event models, we predicted the rate of excursions and of emigration using metrics of soaring flight performance or of daily movement. Contrary to our prediction, we found that soaring performance failed to predict the rate of either excursions or emigration, whereas daily movement predicted both rates, but inconsistently. This was likely because those coarser metrics captured motivation and weather in addition to flight skills. Thus external factors seem more likely to explain the timing of independence and internal development—in this case of flight skills—may be completed quickly. Juveniles may delay emigration not due to lack of skill, but because the costs of leaving parental care outweigh the benefits. The difference we discovered between when individuals were capable of independence and when they committed to it may obscure the more general link between skill acquisition and the end of parental care in other systems as well.

## INTRODUCTION

Independence from parental care is an important life history milestone for many animals. The duration of the dependence period is directly linked to the level of parental investment, which in turn affects the lifelong fitness of both parents and offspring. Parents must balance the trade-off between investing in their current offspring to maximize their survival post-independence and the potential costs this investment may have on the parent’s future reproduction and survival [1–3]. While in some species independence, defined as the ability to operate without parental provisioning and care, can correspond to achieving adult-like body size, for species that occupy complex niches, several key abilities are prerequisite to independence [4, 5]. Particularly, to be independent individuals must have sufficient foraging skill to locate, access, and perhaps process enough nutritious food [6–10]. Additionally, individuals must be capable of avoiding predation and competition without parental protection, which requires them to find safety and compete for opportunities independently [11–13].

Because independent individuals must have these prerequisite abilities, determining their rate of acquisition can allow us to predict the transition to independence. First, individuals must develop physiologically—for example, gain the lung capacity to dive for prey [14], grow feathers that support flight, or build the musculoskeletal structures to accelerate and escape predators [15]. Second, animals must reach behavioral maturity through foraging, fighting, and refined movement skills. For many animals, these skills may be highly specialized—for example, cryptic prey approaching [16] or manipulation of tools [17]. Finally, individuals require environments that support their bids for independence including parental tolerance while their skills develop [18–22], available food [23, 24], suitable environments [25, 26], and potentially either social partners or sufficiently low competition [27–29]. These physical, behavioral, and habitat-specific criteria having been met, individuals can depart parental care.

Intuitively, independence is thought of as the timing of fledging in birds [30] and weaning in mammals [31]. However, for species that require post-fledging and post-weaning provisioning by the parents, initiation of dispersal from the natal areas (hereafter emigration) has been used as an indication of independence [21, 32, 33]. Yet in some systems, adult-level skill competence is reached before emigration [34–41]. For example, although New Caledonian Crows need 10-12 months to achieve adult-level proficiency in tool use, they do not emigrate until around 20 months of age [42]. This raises the question of why the timing of emigration decouples from the rate of skill acquisition, resulting in extended parental care.

In many systems with extended parental care, emigration is preceded by excursions from the natal area (also termed forays or simply extra-territorial movements). These excursions serve as prospecting opportunities [43], providing information about the environment in which dispersal will take place. Animals on excursions can acquire information about competitors, habitat quality, and mate availability, allowing increased post-dispersal breeding success [44]. Excursion rates are often associated with features of the natal territory such as size [45] or resource availability [46] or features of the individual such as sex or social status [45, 47]. While on excursions, individuals are not under parental care or protection and must operate independently. This can be accompanied by stress [48], but potentially delays the dangers associated with dispersal while allowing information gathering—a strategy dubbed stay- and-foray [49]. Thus periods of extended parental care can be punctuated by independent excursions dependent on skill acquisition.

Emigration timing may be influenced by factors other than skill, perhaps including food availability [50, 51] and competitive ability [52]. Importantly, the decision of when to emigrate is not at the sole discretion of the juvenile, but indicates parental motivation either to continue providing care or to begin a new breeding effort.

Here we explore the role of skill acquisition in determining the timing of independence in the context of movement-related skills. We suggest that, at least for movement-related skills, excursions before dispersal can be a better indicator of sufficient skill level for independence than is the decision to emigrate. We test this by investigating the link between how quickly individuals acquire movement capacity and how quickly they begin excursions or emigrate from their parents’ territory. We expect that animals that quickly acquire movement capacity will go on their first excursions relatively early, but that this will not predict early emigration because skilled individuals can still acquire other benefits from parental care.

We used high-resolution GPS-tracking data to explore this in the context of flight skill acquisition in the golden eagle (*Aquila chrysaetos*), a long-lived species that has an extended period of parental care. Golden eagles are obligate soaring birds with high individual- and population-specific variation in the post-fledging dependence period. The duration of the post-fledging dependence period varies from roughly 32 to 260 days in the Western United States [53, p. 666], 39 to 250 days in Scotland [54], 44 to 395 days in France [55], 63 to 176 days in Norway [56], 84 to 269 days in Greece [57], and 143-283 days in Spain [53, pp. 200, 237]. Soaring flight is a vital component of the movement repertoire of golden eagles, allowing them to move the long distances required to hunt, scavenge, and disperse while incurring very low energetic costs. Thus, we can expect learning to soar to predict when they are capable of independence. However, improvements in golden eagle soaring flight performance have been shown to cease well before emigration [22, 55].

We tested two alternative explanations for the lack of relationship between flight performance and emigration: (i) proxies quantifying movement capacity have been too coarse to capture the actual development of movement skill [58]. Previous studies used daily movement metrics (e.g. speed and distance) [22, 55], whereas soaring flight occurs over smaller timescales of minutes. We argue that using fine scale soaring performance may reveal patterns of skill acquisition that coarse scale daily movement metrics do not because soaring performance should be less influenced by motivation regarding whether or not to move. (ii) Emigration timing is not a reliable indicator of behavioral independence as it may be influenced by multiple factors, of which skill proficiency is only one.

## METHODS

### Data collection

We used biologging data from a long-term study of juvenile golden eagles in the Alps. Juvenile eagles were tagged in the nest at approximately 50 days post-hatching (determined using plumage) using solar-powered GPS transmitters from e-obs GmbH (Munich, Germany) and a leg-loop harness [59]. The mass of the tag and harness together was 60 g, constituting only 1.8% of the average mass at tagging for these eagles (mean = 3.42 kg, minimum = 2.06 kg). We used data collected for a total of 97 juveniles over 2019-2024 (downloaded from the Movebank repository on January 25, 2025). GPS data were recorded every 20 min. Additionally, the GPS loggers collected “super bursts” when the batteries were charged above a threshold, which mostly occurred during mid-day. These bursts included 15 minute collection of 1 Hz GPS. We used the super bursts for our analysis of soaring flight before emigration.

### Identification of emigration and excursions

Emigration is defined as leaving the natal territory and not returning at all or returning only briefly, marking the start of natal dispersal. Determining emigration dates can be challenging, especially for excursive species [54]. But for golden eagles, the area used by juveniles is a good indicator of the territory that their parents defend [60]. We manually drew polygons around the highest density of early locations for each individual (e.g. Figure S1) and defined this as the natal territory. For territories inside of Switzerland, we compared our polygons to those published in Jenny et al. [61, p. 368] and found they were in agreement, but we had no such comparisons for other countries. We defined emigration date as the day that an eagle in its first year of life left its natal territory and either did not return at all or returned for less than 24 hours. Young eagles that stopped transmitting within 10 days of leaving the territory (n = 4) were considered inconclusive and we did not count these as emigrations. We defined excursions as any movement outside of the natal territory before emigration, and focused our analysis on long-distance (≥ 10 km) excursions (see supplemental code).

### Classification of flight modes

We identified soaring and gliding flight using the 1 Hz GPS data before emigration. We isolated bursts of continuous 1 Hz GPS data and calculated turning angle and vertical speed between consecutive locations (using the *move* R package [62]). For the classification of the flight behavior, we applied a moving window of 25 s to calculate the absolute cumulative sum of the turning angles (hereafter cumulative turning angle) and a moving window of 5 s to calculate the average vertical speed. We then applied a *k* -means clustering algorithm with *k* = 2 on the average vertical speed to distinguish between soaring and gliding behavior. We sub-categorized soaring observations into circular soaring or linear soaring based on the cumulative turning angle, where a value ≥ 200 degrees indicated circular soaring. Observations with a ground speed ≤ 3.5 ms^−1^ were considered as non-classified (“other behavior”).

### Soaring performance metrics

We categorized each bout of circular soaring lasting at least 30 seconds and separated by not more than 60 seconds as a thermal. For each thermal, we calculated: (i) the vertical speed (height gained per time spent), (ii) wind-corrected circling radius (using the *moveWindSpeed R* package [63, 64]), and (iii) the mean of the 1 Hz wind speeds [64]. We expected greater vertical speeds to reflect better exploitation of thermals, smaller circling radii to indicate soaring closer to the thermal core (where thermals are strongest), and higher wind speeds to indicate more skillful use of thermals as wind can shear and move uplifts.

### Daily movement metrics

We extracted four metrics describing how young golden eagles moved on a daily basis. To consider overall mobility, we estimated ground speeds by down-sampling the data to consistent (20 minute) intervals and extracting the daily maximum speed. On the assumption that climbing in thermals requires skill, we extracted the maximum height above ellipsoid per day to reflect how high individuals flew [55]. We estimated the proportion of time spent in flight as observations per day with ground speeds ≥ 3.5 m/s given all observations on that day [55]. Finally, we down-sampled the data to one location in the hour of noon or 1:00 pm and calculated distances between consecutive midday locations [22] (with the Vincenty method using the *geosphere R* package [65]).

### Time-to-event analysis

To understand the factors that contributed to timing of emigration and the first long-distance excursion, we built four separate time-to-event models (Table 1). The models predicting emigration used only data up to and including the date of emigration (i.e., the post-fledging dependence period) and the models predicting first long-distance excursions used only data up to and including the first day of that excursion. The models used intervals of one day, but our soaring performance metrics were measured at the per-thermal level. Thus, we summarized our soaring performance data to per-day medians of circling radius, vertical speed, and wind speed. We coded each day with a binary outcome (event = 1, no event = 0), with the day right-censored (event = 0) if the event (either first long-distance excursion or emigration) was not observed. The fine-scale data had gaps on days when the biologging devices did not transmit data classified as thermal soaring. If there were no data on the day of excursion/emigration, but there were data within ten days after this (excursions n = 3, emigration n = 6), we used the first of those thermals for the final skill level (i.e. last follow-up time). For the daily-scale metrics, the final skill level was measured on the day of first excursion or emigration.

**Table 1.**
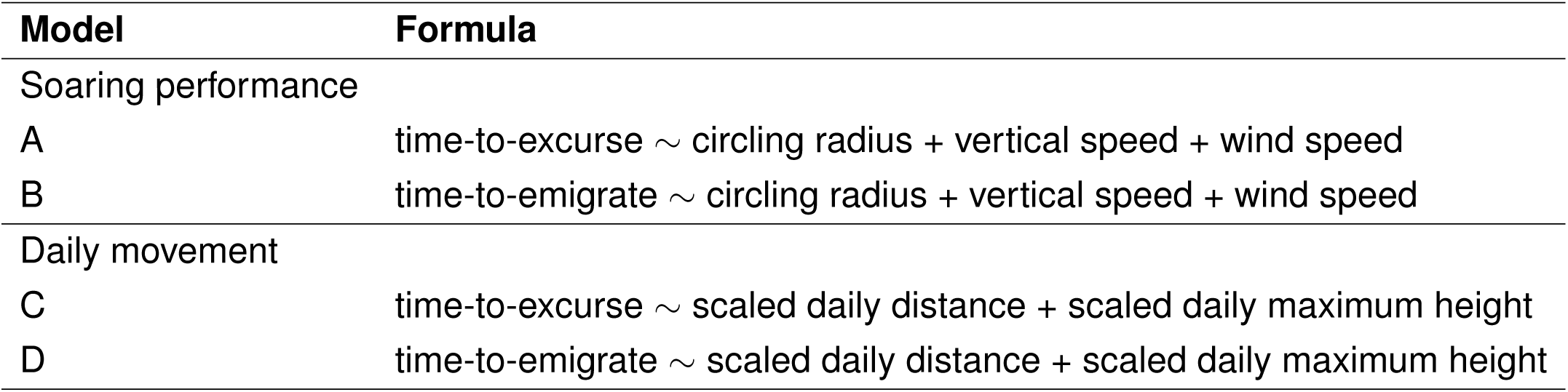
The four time-to-event models using movement metrics to predict latency to first excursion or to emigration. We predicted each event (first long-distance excursion or emigration) at two scales: i) within soaring performance metrics summarized to per-day medians, and ii) the daily movements.

Daily distance traveled between midday locations was correlated with maximum daily ground speed (r = 0.50) and time spent in flight (r = 0.55), thus we only included daily distance and maximum flight height (which had a correlation of r = 0.17) in models C and D (Table 1). Because golden eagles can fly at up to 30 m/s [66], we z-scaled the daily movement metrics to use units of standard deviations rather than individual meters. We built separate models for the daily movement and soaring performance predicting time-to-event using Cox proportional hazards in the *survival* [67] package in *R*. For all four models, we checked the proportional hazards assumption and the distribution of the Schoenfeld residuals [68].

## RESULTS

Of the 97 juveniles, 65 emigrated before the spring of 2025 and 63 performed long-distance excursions. Only 14 individuals transmitted 1 Hz data containing classified thermal soaring flight before going on a long-distance (≥ 10 km) excursion and 18 before emigration. These subsets of 14 and 18 individuals contributed data to our analysis of soaring performance metrics (Figure 1 A-C). For the daily movement metrics (Figure 1 D & E), 65 individuals contributed to the emigration models and 63 to the excursion models.

**Figure 1.**
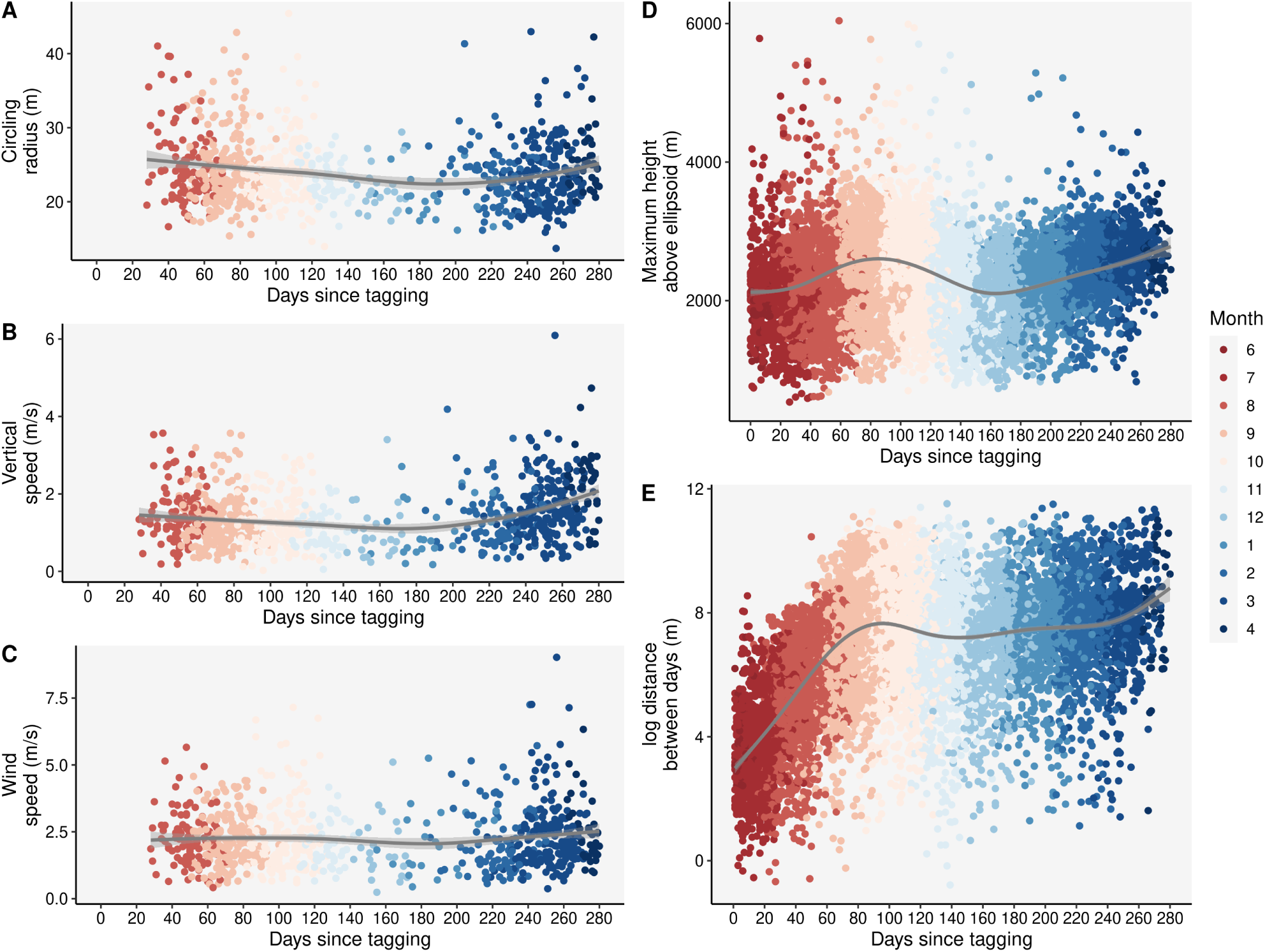
The data used in the models from Table 1. A) Circling radius within thermal uplifts varies and decreases slightly after 120 days. B) Vertical speeds of thermals (meters gained in a thermal per seconds spent in that thermal) vary. C) Wind speeds experienced within thermals vary. D) Maximum height reached per day initially increases until around 80 days after tagging, but then decreases until 160 days before increasing again. E) Distances between consecutive midday locations increases until around 80 days after tagging and then levels off. The median days-to-excursion was 79. The interval of 90 to 160 days corresponds roughly to October through December.

Eagles made their first long-distance excursions between 55 days and 246 days after tagging (median = 79, Figure S2 A). The maximum distances reached on these excursions varied between 10 km and 77 km (median = 16.7, Figure S2 B) and lasted between one hour and 13 days (median = 4.82 hours, Figure S2 C). Emigration occurred between 61 days and 283 days after tagging (median = 229, Figure S2 D).

None of the soaring performance metrics was significantly associated with excursion or emigration rates (Figure 2 A & B, Figure S3 A & B). However, the daily movement metrics had effects on both time-to-excursion and time-to-emigration (Figure 2 C & D, Figure S3 C & D). For each standard deviation increase in daily distance (one sd = 6736 m) and in maximum daily height (one sd = 558 m), we found corresponding increases in the rate of excursions of 264% (hazard ratio = 3.64) and 86% (hazard ratio = 1.86) respectively, thus individuals traveling farther and flying higher were more likely to go on their first excursions (Table 2). Each standard deviation increase in daily distance was associated with a 14% (hazard ratio = 1.14) increase in rate of emigration (Table 2). We found no significant effect of maximum daily height on rate of emigration. We did find a seasonal pattern of lower daily heights and daily distances in winter (Figure 1).

**Figure 2.**
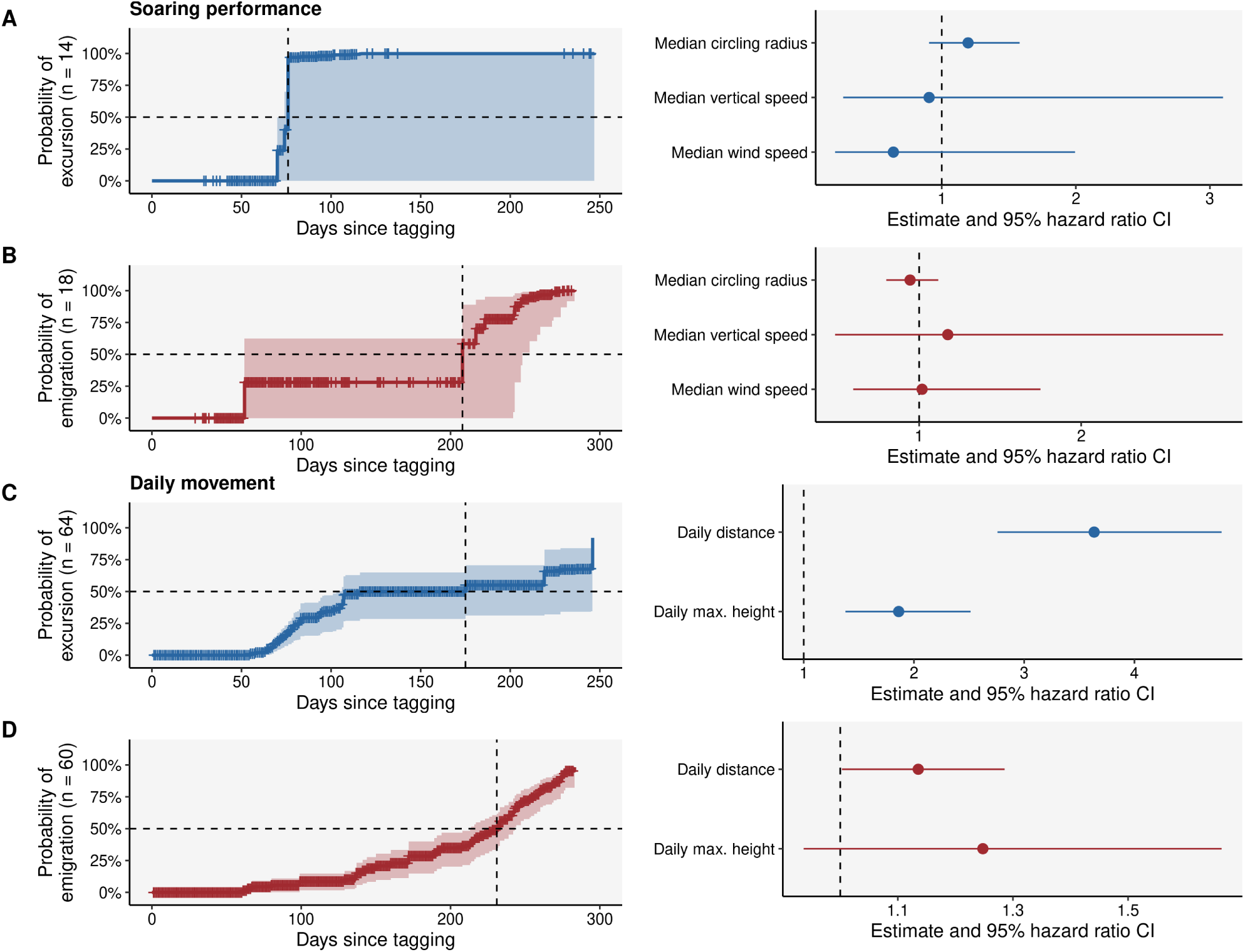
The association of soaring performance and daily movement metrics with time-to-excursion and time-to-emigrate. A) No soaring performance metric had a detectable effect on excursion timing. B) No soaring performance metric had a detectable effect on emigration timing. C) Both daily distance and daily maximum flight height had positive effects on time-to-excursion. D) Daily distance had a positive effect and maximum flight height had no significant effect on time-to-emigrate.

**Table 2.** Hazard ratios and associated confidence from cox proportional hazards tests. Models A and C predicted time-to-excursion, models B and D predicted time-to-emigration (see Table 1).

| Soaring performance |  |  |  |  |  |
| --- | --- | --- | --- | --- | --- |
| Model | Predictor | Hazard Ratio | 2.5% CI | 95% CI | p-value |
| A | Circling radius | 1.197 | 0.906 | 1.581 | 0.207 |
| A | Vertical speed | 0.906 | 0.265 | 3.098 | 0.875 |
| A | Wind speed | 0.640 | 0.205 | 1.994 | 0.441 |
| B | Circling radius | 0.944 | 0.797 | 1.119 | 0.507 |
| B | Vertical speed | 1.177 | 0.482 | 2.876 | 0.721 |
| B | Wind speed | 1.018 | 0.593 | 1.749 | 0.947 |
| Daily movement |  |  |  |  |  |
| Model | Predictor | Hazard Ratio | 2.5% | 95% | p-value |
| C | Daily distance | 3.635 | 3.757 | 4.790 | 5.18e-20 |
| C | Daily maximum height | 1.862 | 1.379 | 2.514 | 5.046e-5 |
| D | Daily distance | 1.135 | 1.003 | 1.286 | 0.045 |
| D | Daily maximum height | 1.248 | 0.936 | 1.663 | 0.131 |

## DISCUSSION

The timing of the transition to independence appeared to depend more on external conditions than on movement capacity. Traveling longer distances between days and achieving higher flight heights were associated with shorter time-to-excursion, but none of our measures of soaring performance was. Similarly, soaring performance failed to predict time-to-emigrate, although longer daily distances were associated with shorter time-to emigrate. This is likely because young eagles learned to soar within a few months, but daily movement captures more than just movement capacity.

Movement at the daily scale is probably strongly influenced by motivation and environmental conditions. Essentially, even while skills remain the same, motivations may change. For example, whether a carcass is available nearby or whether it is foggy should be strong motivators affecting how far young golden eagles move. An eagle may move greater distances when foraging than when satiated but this may not be matched by a change in its fine-scale soaring performance. In fact, the inconsistent signal we found in daily maximum height suggests an effect of weather rather than skill—eagles are more likely to make their first excursions before winter when they are flying higher, but to emigrate after winter and the corresponding reduction in flight height (Figure 1, Figure S2). Thus, it seems that in analyzing these daily movements, we were not directly addressing movement capacity. Instead, the links we found between daily movement and time-to-excursion or time-to-emigrate could be better indicators of a connection to motivation and environment.

We did not find an association between soaring flight performance and time-to-event, which could be because (i) eagles quickly reach their maximum soaring proficiency, and/or (ii) there was a pattern that our data did not capture. It is possible that the way eagles soar in thermals does not change with age as much as thermal structure itself changes with hour and season. Observations of juvenile eagles flying with their parents suggest that adult-like vertical speeds can be obtained within 60 days of leaving the nest [69, p. 227]. It may be that after approximately two months juveniles are exploiting the thermals they can find at peak proficiency. But we might also not have found a signature of learning because the early learning period of up to 60 days post-fledging accounts for only around 5% of our 1 Hz data. The poor coverage of the very early periods of post-fledging is because nestlings and fledglings tend to sit in the shade, thus the GPS devices rarely had full battery before eagles were spending time soaring with their backs exposed to sunlight. In addition, fledglings simply spend less time soaring (which is evident from behavioral classifications in the acceleration data [70]) and have shorter flight intervals, meaning that we were less likely to capture their flight in our burst data structure. Even if young eagles continue to improve after this early phase, our data were skewed to include more transmissions in the afternoon, which yield records of soaring performance predominantly during good soaring conditions. Thus juveniles could have performed relatively well during the data transmission even if they were not yet proficient in less supportive thermal conditions [71]. Our segmentation approach relied on identifying occasions of circling flight, and thus if juveniles were failing to circle consistently due to their inexperience or lack of skills, we might simply have removed those times and instances when juveniles struggled even if they were included in data transmission. Finally, we only accounted for circular, thermal soaring, but young eagles may rely more on orographic soaring [72] and how they perform in orographic uplifts may prove more important for them than thermals.

Regardless of the methodological shortcomings, before emigration from the natal territory most juvenile eagles were clearly capable of remaining outside of parental care for several days at a time and traversing large distances quickly, as demonstrated by their excursions. In addition, four young eagles departed their natal territories in 80 days or fewer. Although our sample was not yet large enough for a formal test, these early departures suggest that movement capacity may not be a constraint on emigration after 80 days. This time-frame is close to the age at which most Alpine eagles started their long-distance excursions (Figure S2) and to the ages at which Alaskan (63 days [73]) and Norwegian (100 days [56]) golden eagles migrate. Eagle parents may stop actively provisioning juveniles at around 120 days [69, p. 228], which could also indicate that juveniles are capable of scavenging for themselves within four months of leaving the nest (although, due to snow, this task becomes more difficult once winter sets in). Taken together, this suggests that golden eagles achieve the ability to be independent with regard to maturity of movement capacity in three to four months. Thus, in contrast to the hypothesis that acquisition of complex skills delays emigration, it seems that soaring performance is unlikely to act as a constraint on eagle emigration after a very early age. In addition, this ability to be independent is better captured by excursion timing than by emigration timing.

Our findings suggest that the transition to independence in juvenile golden eagles is not a simple process following the smooth trajectory of a learning curve, but shaped by a combination of development, environment, and decisions. The variability in golden eagle emigration highlights that independence is often a choice. For golden eagles, soaring is a vital skill that allows them to hunt and scavenge, but they seem to acquire this skill—at least under optimal soaring conditions—long before they emigrate. Instead, they may engage in long-distance excursions after they acquire adult-like movement abilities, using excursions as an opportunity to practice moving and scavenging in unfamiliar environments and to prospect for available habitats from the safety of their natal territories [22, 56, 74, 75]. The golden eagle population in the Alps is at its carrying capacity, with no territories unoccupied [61, p. 370]. Thus juveniles that are otherwise ready to disperse may simply have nowhere to go. Competitive ability, parental tolerance, conspecific densities, prey densities, hunting skills, and human disturbance are among many factors both internal and external to a young animal that are likely to interact in affecting the decision of whether or not to forgo parental care. Whereas movement capacity is certainly a necessary part of the ability to survive independently, it is just one factor contributing to the complex decision of whether and when to become independent.

The transition to independence is often associated with mortality. Unless the expected profit of undertaking this risky transition is great enough, young animals that can exploit parental tolerance should continue to do so, even if they could potentially survive on their own. Both the costs and the benefits of leaving parental care are probably best assessed during excursions from the natal territory, which may link excursion timing to when individuals have reached the developmental stage to attempt independence. Although movement capacity may be acquired relatively early in development, emigration often occurs later, which indicates that the timing of independence can be decoupled from skill acquisition. Delaying independence allows juveniles to choose the timing of emigration, aligning it with favorable external conditions rather than with internal developmental milestones. Thus juveniles may remain in the natal area not because they are unskilled, but because the costs of leaving parental care outweigh the benefits.

## ETHICS

The handling and ringing of golden eagle nestlings in Switzerland was carried out under the authorization of the Office for Food Safety and Animal Health (ALT) of the canton Grisons (licence No. GR 2017 06, GR 2018 05E, GR 2019 03E). In Italy, the permissions for handling, tagging and marking were obtained from autonomous region of South Tyrol (Dekret 12257/2018 and Dekret 8788/2020), as well as from the region of Lombardia for ringing and tagging through in Lombardia and South Tyrol by ISPRA (Istituto Superiore per la Protezione e la Ricerca Ambientale) with the Richiesta di autorizzazione alla cattura di fauna selvatica per scopi scientifici(l.r. 26/93). In Austria, all procedures for handling, tagging and marking were approved by the Ethics committee of the University of Vienna (No. 2020-008) and permitted by the Federal Ministry for Education, Science and Research (No. 2020-0.547.571), Styria (BHLI-165942/2021-2) and Upper Austria (LFW-2021-263262/7-Sr). Finally, in Germany birds handled, tagged and ringed were done so under the permission issued by the government of Oberbayern (2532.Vet 02-16-88 and 2532.Vet 02-20-86). All procedures followed the ASAB/ABS guidelines for the ethical treatment of animals in behavioral research and teaching and all applicable international, national, and/or institutional guidelines for the care and use of animals were followed. The handling of birds was performed with maximum care and minimal disturbance to nests and the landscape. Ethical approval for involving animals in this study was received through the application procedure for ringing permits and the scientific commission of the Swiss Ornithological Institute and the national authorities as well as the guidelines imposed through the European Commission on the basis of the guidelines of the FELASA (the Federation or the European Laboratory Animals Association).

## DATA ACCESSIBILITY

The raw tracking data used for this study will be made available via Movebank upon acceptance of the manuscript.

## DECLARATION OF AI USE

We have not used AI-assisted technologies in creating this article.

## AUTHORS’ CONTRIBUTIONS

EN, KS, and HB conceived of the study. KS, EB, WF, JSH, PS, MT, and MW facilitated the fieldwork and data collection. MC assisted with time-to-event analysis. SZ provided initial estimations of emigration timing. HB and MS processed the data. HB analyzed the data and wrote the first draft of the manuscript. All authors discussed the final results and commented on the manuscript.

## ACKNOWLEDGMENTS.

We are grateful to Emily Shepard for discussions on estimation of soaring skills metrics. We are sincerely grateful for the extensive efforts involved in tagging young eagles by all those listed in Nourani et al. 2024 [72]. We thank Andrea Flack for her patient support.

## SUPPLEMENTARY INFORMATION

**Table S1.** The definitions of important terms used in this paper along with the page and paragraph number of their first use or definition in the text.

| Term | Meaning | Page, paragraph |
| --- | --- | --- |
| Skill acquisition | obtaining new abilities and/or improving existing abilities | 2, 2 |
| Fledging | leaving the nest permanently | 3, 2 |
| Emigration | leaving the natal territory permanently; the start of natal dispersal | 3, 2 |
| Excursion | extra-territorial movements preceding emigration | 3, 3 |
| Soaring performance | how successfully individuals use soaring flight (e.g. vertical speed) | 4, 2 |
| Daily movement | how individuals move at the daily scale (e.g. total distance traveled) | 4, 3 |
| Vertical speed | height gained in a thermal per time spent | 6, 2 |
| Circling radius | the radius of the circle a soaring bird has to turn to remain inside a thermal column | 6, 2 |
| Wind speed | high speed winds can shear uplifts and make them more difficult to ride | 6, 2 |
| Daily distance | displacement distance between midday observations | 6, 3 |
| Daily max. height | maximum height above the ellipsoid attained in a day | 6, 3 |

**Figure S1.**
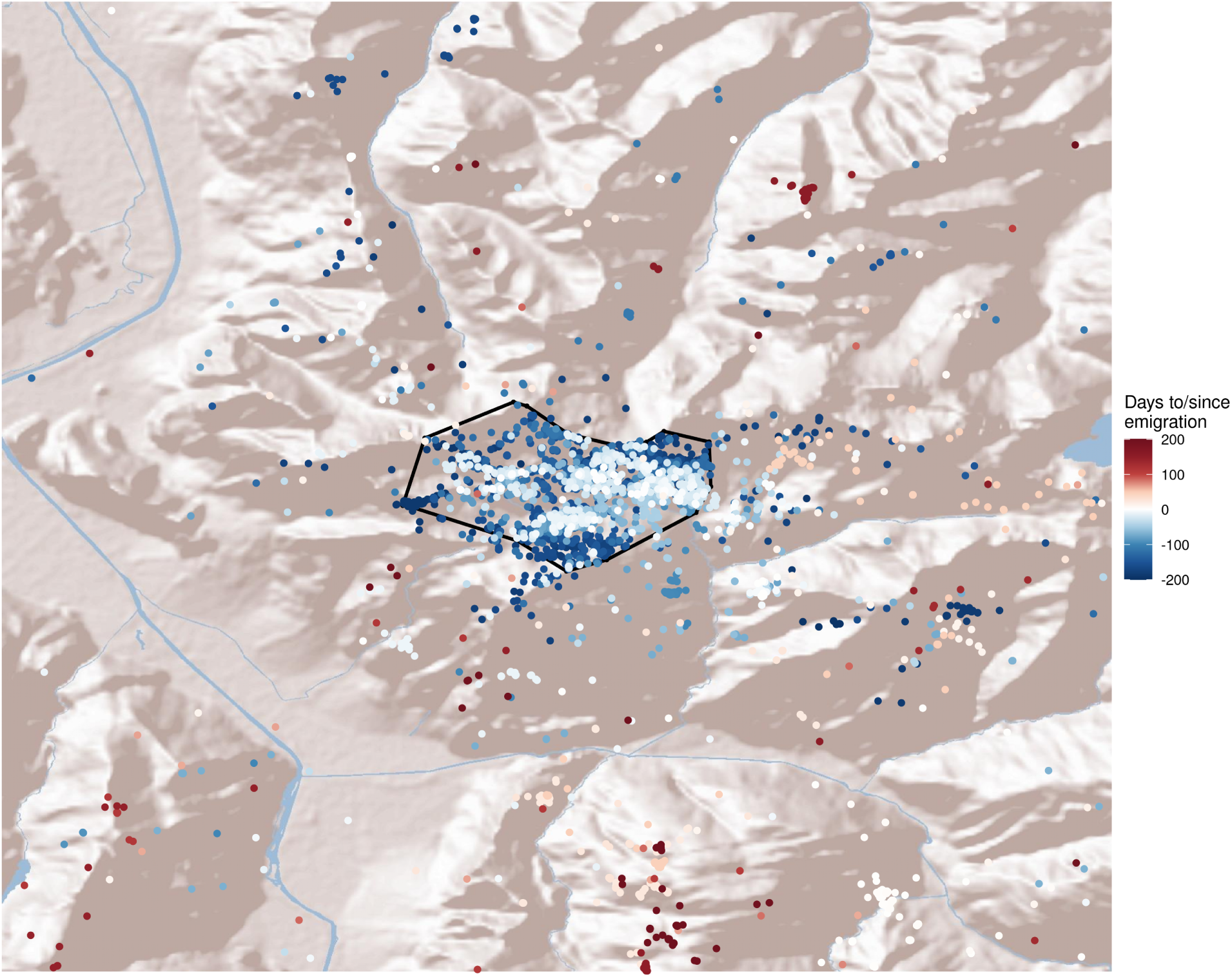
An example of a natal territory polygon in the context of terrain (OpenStreetMap data). Points in blue are pre-dispersal, white points are from the emigration date, and red points are during dispersal.

**Figure S2.**
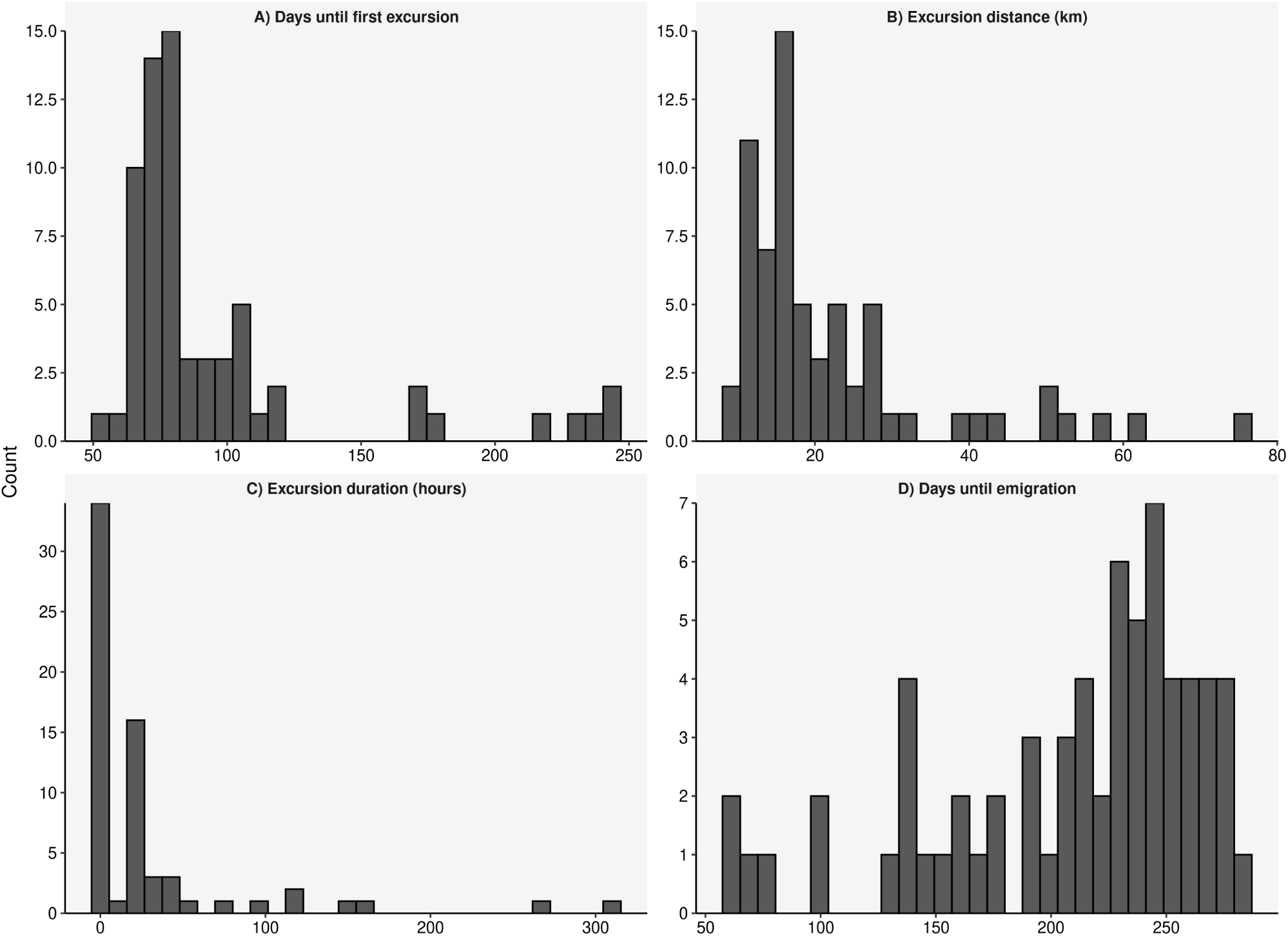
A) 75% of young golden eagles had made their first long-distance excursion within 100 days of tagging (roughly 145-155 days since hatching). B) These first excursions tended to be within 26 km of the natal territory border. C) These first excursions had a bimodal distribution of durations with one peak between 1 and 6 hours and another between 16 and 30 hours. D) Almost two thirds of the eagles emigrated after 200 days (n = 45) and only one third emigrated earlier (n = 21).

**Figure S3.**
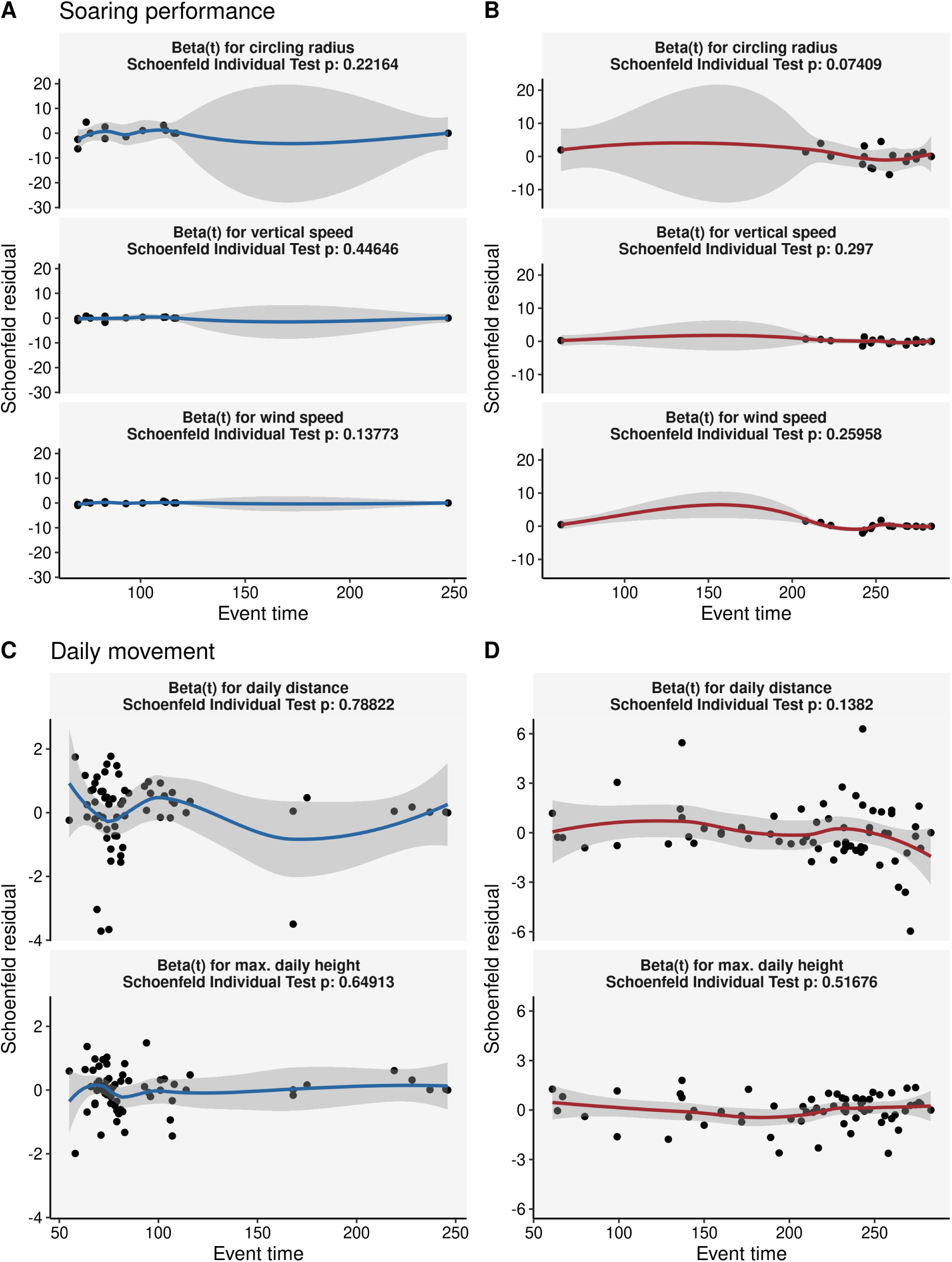
The Schoenfeld residuals show no significant time-dependent effects. A) The residuals for fine-scale data predicting time-to-excursion. B) The residuals for fine-scale data predicting time-to-emigration. C) The residuals for daily-scale data predicting time-to-excursion. D) The residuals for daily-scale data predicting time-to-emigration. These residuals correspond to panels A-D of Figure 2.

## REFERENCES

1. Johnson, L. S., Nguyen, A. V. & Connor, C. L. Do parents play a role in the timing and process of fledging by nestling Mountain Bluebirds? Journal of Field Ornithology 88, 39–46 (2017).

2. López-Idiáquez, D., Vergara, P., Fargallo, J. A. & Martínez-Padilla, J. Providing longer post-fledging periods increases offspring survival at the expense of future fecundity. PLOS ONE 13, 1–13. 10.1371/journal.pone.0203152 (Sept. 2018).

3. Jones, T. M. et al. Parental benefits and offspring costs reflect parent–offspring conflict over the age of fledging among songbirds. Proceedings of the National Academy of Sciences 117, 30539–30546. eprint: https://www.pnas.org/doi/pdf/10.1073/pnas.2008955117. https://www.pnas.org/doi/abs/10.1073/pnas.2008955117 (2020).

4. Charnov, E. L. Evolution of mammal life histories. Evolutionary Ecology Research 3, 521–535 (2001).

5. Schuppli, C., Isler, K. & van Schaik, C. P. How to explain the unusually late age at skill competence among humans. Journal of human evolution 63, 843–850 (2012).

6. Meinertzhagen, R. The education of young ospreys. Ibis 96, 153–155 (1954).

7. Davies, N. & Green, R. The development and ecological significance of feeding techniques in the Reed Warbler (Acrocephalus scirpaceus). Animal Behaviour 24, 213–229 (1976).

8. Kitowski, I. Play behaviour and active training of Montagu’s harrier (Circus pygargus) offspring in the post-fledging period. Journal of Ethology 23, 3–8 (2005).

9. Catlin, D. H., Felio, J. H. & Fraser, J. D. Effects of water discharge on fledging time, growth, and survival of piping plovers on the Missouri River. The Journal of wildlife management 77, 525–533 (2013).

10. Geipel, I., Kalko, E. K., Wallmeyer, K. & Knörnschild, M. Postweaning maternal food provisioning in a bat with a complex hunting strategy. Animal Behaviour 85, 1435–1441 (2013).

11. Gersick, A. S., Snyder-Mackler, N. & White, D. J. Ontogeny of social skills: social complexity improves mating and competitive strategies in male brown-headed cowbirds. Animal Behaviour 83, 1171–1177 (2012).

12. Putman, B. J., Coss, R. G. & Clark, R. W. The ontogeny of antipredator behavior: age differences in California ground squirrels (Otospermophilus beecheyi) at multiple stages of rattlesnake encounters. Behavioral Ecology and Sociobiology 69, 1447–1457 (2015).

13. York, C. A. & Bartol, I. K. Anti-predator behavior of squid throughout ontogeny. Journal of Experimental Marine Biology and Ecology 480, 26–35 (2016).

14. Grecian, W. J. et al. Environmental drivers of population-level variation in the migratory and diving ontogeny of an Arctic top predator. Royal Society Open Science 9, 211042 (2022).

15. Herrel, A. & Gibb, A. C. Ontogeny of performance in vertebrates. Physiological and biochemical Zoology 79, 1–6 (2006).

16. Schiel, N., Souto, A., Huber, L. & Bezerra, B. M. Hunting strategies in wild common marmosets are prey and age dependent. American journal of primatology 72, 1039– 1046 (2010).

17. Rutz, C. et al. The ecological significance of tool use in New Caledonian crows. Science 329, 1523–1526 (2010).

18. Heinsohn, R. G. Slow learning of foraging skills and extended parental care in co-operatively breeding white-winged choughs. The American Naturalist 137, 864–881 (1991).

19. Castillo-Guerrero, J. A. & Mellink, E. Maximum diving depth in fledging blue-footed boobies: Skill development and transition to independence. The Wilson Journal of Ornithology 118, 527–531 (2006).

20. Ridley, A. R. & Raihani, N. J. Variable postfledging care in a cooperative bird: causes and consequences. Behavioral Ecology 18, 994–1000 (2007).

21. Davies, N. Parental care and the transition to independent feeding in the young spotted flycatcher (Muscicapa striata). Behaviour, 280–295 (1976).

22. Weston, E. D., Whitfield, D. P., Travis, J. M. & Lambin, X. The contribution of flight capability to the post-fledging dependence period of golden eagles. Journal of Avian Biology 49 (2018).

23. Engler, M. & Krone, O. Movement patterns of the White-tailed Sea Eagle (Haliaeetus albicilla): Post-fledging behaviour, natal dispersal onset and the role of the natal environment. Ibis 164, 188–201 (2022).

24. Scherler, P. et al. Determinants of departure to natal dispersal across an elevational gradient in a long-lived raptor species. Ecology and Evolution 13, e9603 (2023).

25. Molteno, A. & Bennett, N. Rainfall, dispersal and reproductive inhibition in eusocial Damaraland mole-rats (Cryptomys damarensis). Journal of Zoology 256, 445–448 (2002).

26. Walls, S. S., Kenward, R. E. & Holloway, G. J. Weather to disperse? Evidence that climatic conditions influence vertebrate dispersal. Journal of Animal Ecology, 190–197 (2005).

27. Nicolaus, M. et al. Social environment affects juvenile dispersal in great tits (Parus major). Journal of Animal Ecology 81, 827–837 (2012).

28. Perry, S., Godoy, I., Lammers, W. & Lin, A. Impact of personality traits and early life experience on timing of emigration and rise to alpha male status for wild male white-faced capuchin monkeys (Cebus capucinus) at Lomas Barbudal Biological Reserve, Costa Rica. Behaviour 154, 195–226 (2017).

29. Sen, S. et al. Social correlates of androgen levels and dispersal age in juvenile male geladas. Hormones and Behavior 146, 105264 (2022).

30. Mainwaring, M. C. The transition from dependence to independence in birds. Behavioral ecology and sociobiology 70, 1419–1431 (2016).

31. Lee, P. C. The meanings of weaning: growth, lactation, and life history. *Evolutionary Anthropology: Issues, News, and Reviews: Issues*, News, and Reviews 5, 87–98 (1996).

32. Bustamante, J. Post-fledging dependence period and development of flight and hunting behaviour in the red kite Milvus milvus. Bird Study 40, 181–188 (1993).

33. Jenkins, J. M., Thompson III, F. R. & Faaborg, J. Behavioral development and habitat structure affect postfledging movements of songbirds. The Journal of Wildlife Management 81, 144–153 (2017).

34. Johnson, S. J. Development of hunting and self-sufficiency in juvenile red-tailed hawks (Buteo jamaicensis). Journal of Raptor Research 20, 4 (1986).

35. Donazari, J. A. & Ceballos, O. Post-fledging dependence period and development of flight and foraging behavior in the Egyptian vulture Neophron percnopterus. Ardea 78, 387–394 (1990).

36. Langen, T. A. Skill acquisition and the timing of natal dispersal in the white-throated magpie-jay, Calocitta formosa. Animal Behaviour 51, 575–588 (1996).

37. Wheelwright, N. T. & Templeton, J. J. Development of foraging skills and the transition to independence in juvenile Savannah Sparrows. The Condor 105, 279–287 (2003).

38. Stone, A. I. Foraging ontogeny is not linked to delayed maturation in squirrel monkeys (Saimiri sciureus). Ethology 112, 105–115 (2006).

39. Schuppli, C., Graber, S. M., Isler, K. & van Schaik, C. P. Life history, cognition and the evolution of complex foraging niches. Journal of Human Evolution 92, 91–100 (2016).

40. Collet, J., Prudor, A., Corbeau, A., Mendez, L. & Weimerskirch, H. First explorations: ontogeny of central place foraging directions in two tropical seabirds. Behavioral Ecology 31, 815–825 (2020).

41. Corbeau, A., Prudor, A., Kato, A. & Weimerskirch, H. Development of flight and foraging behaviour in a juvenile seabird with extreme soaring capacities. Journal of Animal Ecology 89, 20–28 (2020).

42. Holzhaider, J. C., Hunt, G. R. & Gray, R. D. Social learning in New Caledonian crows. Learning & Behavior 38, 206–219 (2010).

43. Reed, J. M., Boulinier, T., Danchin, E. & Oring, L. W. Informed dispersal: prospecting by birds for breeding sites. Current ornithology, 189–259 (1999).

44. Pärt, T., Arlt, D., Doligez, B., Low, M. & Qvarnström, A. Prospectors combine social and environmental information to improve habitat selection and breeding success in the subsequent year. Journal of Animal Ecology 80, 1227–1235 (2011).

45. Mayer, M., Zedrosser, A. & Rosell, F. Extra-territorial movements differ between territory holders and subordinates in a large, monogamous rodent. Scientific Reports 7, 15261 (2017).

46. Messier, F. Solitary living and extraterritorial movements of wolves in relation to social status and prey abundance. Canadian Journal of Zoology 63, 239–245 (1985).

47. Kamler, J. F., Stenkewitz, U., Gharajehdaghipour, T. & Macdonald, D. W. Social organization, home ranges, and extraterritorial forays of black-backed jackals. The Journal of Wildlife Management 83, 1800–1808 (2019).

48. Young, A. J. & Monfort, S. L. Stress and the costs of extra-territorial movement in a social carnivore. Biology Letters 5, 439–441 (2009).

49. Walters, J. R., Doerr, P. D. & Carter III, J. Delayed dispersal and reproduction as a life-history tactic in cooperative breeders: fitness calculations from red-cockaded woodpeckers. The American Naturalist 139, 623–643 (1992).

50. Cucherousset, J., Paillisson, J.-M. & Roussel, J.-M. Natal departure timing from spatially varying environments is dependent of individual ontogenetic status. Naturwis-senschaften 100, 761–768 (2013).

51. Fattebert, J., Perrig, M., Naef-Daenzer, B. & Grüebler, M. U. Experimentally disentangling intrinsic and extrinsic drivers of natal dispersal in a nocturnal raptor. Proceedings of the Royal Society B 286, 20191537 (2019).

52. Gjershaug, J. O., Halley, D. & Stokke, B. G. Predefinitive plumage in the golden eagle (Aquila chrysaetos): a signal of aggression or submission? Journal of Raptor Research 53, 431–435 (2019).

53. Bautista-Rodríguez, J. & Ellis, D. The Golden Eagle Around the World ISBN: 978-0-88839-775-1 [Hardback] (Hancock House Publishers, Nov. 2024).

54. Weston, E. D., Whitfield, D. P., Travis, J. M. & Lambin, X. When do young birds disperse? Tests from studies of golden eagles in Scotland. BMC ecology 13, 1–13 (2013).

55. Hemery, A., Mugnier-Lavorel, L., Itty, C., Duriez, O. & Besnard, A. Timing of departure from natal areas by golden eagles is not constrained by acquisition of flight skills. Journal of Avian Biology 2023, e03111 (2023).

56. Nygård, T., Jacobsen, K.-O., Johnsen, T. V. & Systad, G. H. Dispersal and survival of juvenile Golden Eagles (Aquila chrysaetos) from Finnmark, northern Norway. Journal of Raptor Research 50, 144–160 (2016).

57. Sidiropoulos, L. et al. Dispersal Ecology of Golden Eagles (Aquila chrysaetos) in Northern Greece: Onset, Ranging, Temporary and Territorial Settlement. Diversity 16, 580 (2024).

58. Nathan, R. et al. Big-data approaches lead to an increased understanding of the ecology of animal movement. Science 375, eabg1780 (2022).

59. Longarini, A. et al. Effect of harness design for tag attachment on the flight performance of five soaring species. Movement Ecology 11, 1–13 (2023).

60. Hemery, A., Duriez, O., Itty, C., Henry, P.-Y. & Besnard, A. Using juvenile movements as a proxy for adult habitat and space use in long-lived territorial species: a case study on the golden eagle. Journal of Avian Biology, e03212 (2024).

61. Jenny, D. et al. in The Golden Eagle in Switzerland (eds Ellis, D. H., Bautista, J. & Ellis, C. H.) 367–394 (Hancock House Publishers, 2025). ISBN: 978-0-88839-775-1.

62. Kranstauber, B., Smolla, M. & Scharf, A. K. *move: Visualizing and Analyzing Animal Track Data* R package version 4.2.4 (2023). https://CRAN.R-project.org/package=move.

63. Weinzierl, R. et al. Wind estimation based on thermal soaring of birds. Ecology and Evolution 6, 8706–8718 (2016).

64. Kranstauber, B. & Weinzierl, R. *moveWindSpeed: Estimate Wind Speeds from Bird Trajectories* R package version 0.2.4 (2023). https://CRAN.R-project.org/package=moveWindSpeed.

65. Hijmans, R. J. *geosphere: Spherical Trigonometry* R package version 1.5-18 (2022). https://CRAN.R-project.org/package=geosphere.

66. Garstang, M., Greco, S., Emmitt, G. D., Miller, T. A. & Lanzone, M. An instrumented golden eagle’s (aquila chrysaetos) long-distance flight behavior. Animals 12, 1470 (2022).

67. Therneau, T. M. *A Package for Survival Analysis in R* R package version 3.7-0 (2024). https://CRAN.R-project.org/package=survival.

68. Schoenfeld, D. Partial residuals for the proportional hazards regression model. Biometrika 69, 239–241 (1982).

69. Watson, J. The golden eagle 227–228 (Bloomsbury Publishing, 2010).

70. Hatzl, J. S. Tracking behavioural trajectories: early-life effects on natal dispersal patterns in golden eagles (Aquila chrysaetos) PhD thesis (ETH Zurich, 2024).

71. Harel, R., Horvitz, N. & Nathan, R. Adult vultures outperform juveniles in challenging thermal soaring conditions. Scientific reports 6, 27865 (2016).

72. Nourani, E. et al. Developmental stage shapes the realized energy landscape for a flight specialist. eLife 13, RP98818 (2024).

73. McIntyre, C. L. & Collopy, M. W. Postfledging dependence period of migratory golden eagles (Aquila chrysaetos) in Denali National Park and Preserve, Alaska. The Auk 123, 877–884 (2006).

74. Soutullo, A., López-López, P., Cortés, G. D., Urios, V. & Ferrer, M. Exploring juvenile golden eagles’ dispersal movements at two different temporal scales. Ethology Ecology & Evolution 25, 117–128 (2013).

75. Murphy, R. K. et al. First-year dispersal of Golden Eagles from natal areas in the southwestern United States and implications for second-year settling. Journal of Raptor Research 51, 216–233 (2017).

